# The hypertrophic cardiomyopathy mutation G768R makes cardiac myosin a high duty ratio motor

**DOI:** 10.64898/2026.06.02.729673

**Authors:** Divya Pathak, Tugba N. Ozturk, Aminah Dawood, Matthew C. Childers, Michael Regnier, James A. Spudich, Chao Liu, Kathleen M. Ruppel

**Affiliations:** Department of Biochemistry, Stanford University, Stanford, CA, USA; Biosciences and Biotechnology Division, Lawrence Livermore National Laboratory, Livermore, CA, USA; Department of Bioengineering, University of Washington, Seattle, WA, USA

## Abstract

β-cardiac myosin is the primary motor protein in the human heart responsible for force generation by converting chemical energy from ATP hydrolysis to mechanical work. It binds actin, produces force through its powerstroke, and releases actin, and must complete this cycle several times a second within each heartbeat. Like other muscle myosins, cardiac myosin has a low duty ratio of ∼5%, the fraction of time in the actin-bound force-producing state. Here we present a hypertrophic cardiomyopathy (HCM) causing mutation, G768R, that increases cardiac myosin’s duty ratio to >60%, an unprecedented >10x increase unmatched by any previously studied muscle myosin mutation. In recombinantly expressed human β-cardiac myosin subfragment-1 (sS1), G768R dramatically decreases the load-sensitive actin-detachment rate and step size of single molecules measured by optical tweezers. The ∼15x longer actin-bound time combined with only a modest change in overall ATPase rate predicts a duty ratio of >60%. Motility velocity is also severely slowed, as expected given the long bound time and decreased step size. All-atom molecular dynamic simulations of the pre-powerstroke and post-rigor states predict that the mutation alters lever arm priming and reduces ADP pocket opening, providing a possible structural mechanism to corroborate the experimental observations. Finally, ATPase experiments on 2-headed heavy meromyosin (HMM) constructs suggest that G768R destabilizes the autoinhibited state of myosin. Our findings of high duty ratio and reduced autoinhibition provide molecular mechanisms of cardiac hypercontractility and impaired relaxation. A muscle myosin in the heart that stays bound to actin for over half its period is bound to have significant functional and clinical consequences.

## Introduction

Hypertrophic cardiomyopathy (HCM) is the most common inherited heart disorder, affecting up to 1 in 200 individuals globally, and remains the leading cause of sudden cardiac death among young adults and athletes (*1, 2*). Clinically, HCM is characterized by left ventricular hypertrophy, myocyte disarray, interstitial fibrosis, diastolic dysfunction, and a predisposition to arrhythmias. HCM is an autosomal dominant disease of the cardiac sarcomere with a heterogenous clinical presentation, varying widely in age of onset, disease severity, and long-term outcomes (*3–5*). Among the sarcomeric genes implicated in HCM, MYH7, which encodes the β-cardiac myosin heavy chain, is frequently affected (*6–8*). Within MYH7, the converter domain is a critical hotspot for pathogenic variants, associated with higher prevalence of mutations and more severe clinical phenotypes (*9–12*). Here, we present the striking molecular effects of the converter domain HCM mutation G768R.

β-cardiac myosin is a mechanochemical enzyme that functions as the primary force-generating motor in the human ventricle, translating chemical energy from ATP hydrolysis into mechanical work through its cyclic interactions with actin. Over the past decades, dozens of studies from multiple labs have established that HCM-causing mutations in the myosin motor domain can affect any parameter of the ATPase cycle, including overall catalytic rate, force generation and sensitivity, step size, ADP release, and duty ratio (fraction of cycle time that is bound to actin) (reviewed by Spudich (*13, 14*)). Changes to these parameters directly alter the power generated by myosin heads (*15–17*) and thus provide an understanding of the molecular mechanism of disease.

In recent years, our understanding of myosin regulation has expanded beyond simple considerations of catalytic activity and powerstroke mechanics. It is now well appreciated that cardiac myosin, like other myosins (*18–20*), adopts a dynamic equilibrium between active and autoinhibited conformations, the latter being the interacting-heads motif (IHM), wherein myosin heads fold back onto their proximal S2 tail and one another (*21–25*). This autoinhibited structural conformation is thought to correlate with a biochemically defined super-relaxed (SRX) state characterized by markedly reduced ATP turnover (*26–30*). It serves as a critical modulator of cardiac energetics, governing the number of heads available for actin interaction during systole and preserving metabolic efficiency at rest. Disruption of this delicate equilibrium represents another central mechanism in the pathogenesis of HCM complimentary to alterations in motor power generation. Numerous disease-associated mutations, including those both within and remote from the IHM interface, destabilize the autoinhibited state, thereby increasing the proportion of functionally accessible myosin heads and augmenting ensemble force generation (*6, 13, 14, 16, 31–35*).

Within the context of both myosin motor mechanics and IHM, the converter domain is structurally critical. It forms a pivotal coupling region between the catalytic motor domain and the lever arm, transmitting nucleotide-driven conformational changes into lever arm rotation, while also extending the IHM surface. Spatial analyses of HCM mutations showed significant enrichment of disease-associated variants in the converter region (*10*). Clinically, converter domain mutations often manifest early, with increased penetrance, more severe hypertrophy, and poorer prognosis (*9, 10, 36, 37*). Mutations in this region have been associated with both increased and decreased stiffness and force generation at the fiber level (*11, 38–40*), yet unchanged or very subtly altered force generation and kinetics at the molecular level (*41*).

To further understand the effects of HCM-causing mutations in the converter domain, we have analyzed the effects of a previously unstudied missense variant, G768R, located near the distal end of the converter domain of β-cardiac myosin (Fig. 1A). G768R is classified as pathogenic in ClinVar and has been documented in a handful of adult and pediatric HCM patients (*42–47*), including a family with an infant requiring heart transplant (*48*) and a fetus that died one day after birth (*49*). Glycine 768 is highly conserved across myosin heavy chain isoforms (Fig. 1B), and a substitution of the neutral glycine to the basic arginine would be expected to be consequential. As part of the critical converter region immediately preceding the lever arm, G768R is in a position to potentially affect both force transduction and the intermolecular interactions that maintain myosin heads in the sequestered OFF state (IHM/SRX). Thus, we hypothesize that G768R may exert multiple effects on myosin function—altering motor kinetics, such as step size, ATPase activity, load sensitivity, and duty ratio, while simultaneously destabilizing the IHM OFF state.

**Figure 1.**
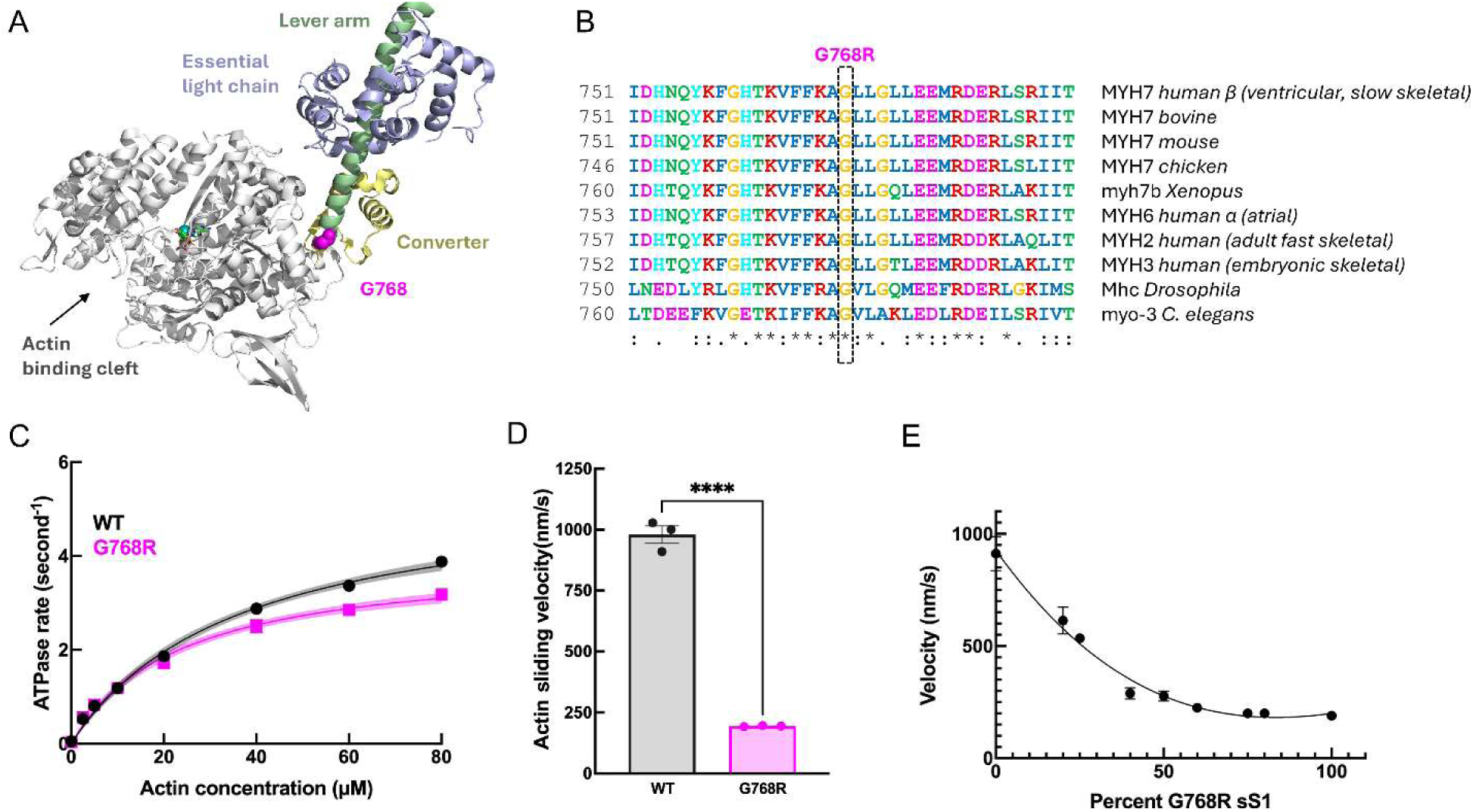
G768R slows motility velocity to a much greater extent than ATPase of the myosin motor domain. **(A)** Location of hypertrophic cardiomyopathy-causing mutation G768R at the end of the converter domain shown as pink spheres on cartoon representation of human β-cardiac myosin sS1 (residues 1-808 of PDB structure 8QYU). **(B)** Multiple sequence alignment of different myosin-IIs expressed in various muscles show that the mutation site is highly conserved. **(C)** Actin-activated ATPase of WT (black) and G768R (pink) sS1. G768R *k*_cat_ = 3.92 ± 0.13 s^−1^,*K*_m_= 25.2 ± 1.2µM, WT *k*_cat_ = 5.37 ± 0.06 s^−1^, *K*_m_= 38.6 ± 2.4µM (see also Fig. S1 and Table S1). **(D)** Actin sliding velocity of WT and G768R sS1 in the unloaded in vitro actin motility assay. Each data point represents a biological replicate. Each biological replicate represents average of three technical replicates. *V*_WT_ = 979.6 ± 61 nm/s, *V*_G768R_ = 194 ± 2 nm/s). Velocities represent MVEL20 from FAST analysis of motility movies (see Methods and Fig. S2). **(E)** Motility velocities for different mixtures of WT and G768R sS1. Error bars are the standard error of mean from at least six movie replicates. Solid line shows the best quadratic fit to the data.

We undertook a comprehensive functional characterization of recombinant human β-cardiac myosin harboring the G768R mutation. Employing an integrated suite of single-molecule biophysical assays, ensemble motility experiments, and steady-state ATPase measurements, we interrogated the impact of G768R on detachment kinetics, duty ratio, powerstroke, and the distribution of myosin heads between relaxed and active states. Unexpectedly, unlike the many other myosin HCM mutations from recent publications whose main effect was disruption of myosin autoinhibition with only minor changes to motor kinetics and mechanics, G768R had its most significant effect on the latter. We found that while it somewhat disrupted the sequestration of myosin heads, G768R dramatically impaired actin detachment and step size on the single molecule level, thereby greatly elevating the duty ratio and reducing actin sliding velocity on the ensemble level. The duty ratio was increased to 0.6, rendering cardiac myosin a high duty ratio motor which has not been observed previously for any mutation or small molecule drug. Finally, we performed atomistic molecular dynamic simulations which suggest that the additional local interactions gained by arginine at position 768 can impede the power stroke and allosterically reduce the opening of the ADP pocket, providing a possible structural mechanism to corroborate the experimental observations.

Collectively, these results suggest that G768R induces hypercontractility through a dual mechanism involving greatly prolonged actin attachment and moderate disruption of myosin’s autoinhibited state. These insights underscore the pivotal and multifaceted role of the converter domain in myosin function and expand our understanding of the molecular mechanisms by which mutations allosterically perturb motor kinetics, mechanics, and ON/OFF equilibrium to drive the pathogenesis of hypertrophic cardiomyopathy.

## Results

### G768R slows motility velocity to a much greater extent than ATPase of the myosin motor domain

The G768R mutation is located at the distal end of the converter domain, immediately preceding the lever arm—a region essential for transducing chemical energy from nucleotide hydrolysis into mechanical movement (Fig. 1A). We first investigated the functional consequences of this mutation on the human β-cardiac myosin motor domain (residues 1-808 of Subfragment-1, sS1). Steady-state actin-activated ATPase assays revealed a small reduction in catalytic activity and increased actin affinity for G768R compared to wildtype (WT) sS1 (G768R *k*_cat_ = 3.92 ± 0.13 s^−1^, *K*_m_= 25.2 ± 1.2 µM; WT *k*_cat_ = 5.37 ± 0.06 s^−1^, *K*_m_= 38.6 ± 2.4 µM; at 23 °C; p<0.05) (Fig. 1C, Fig. S1, Table S1).

By contrast, the mutation showed a dramatically larger effect in the unloaded motility assay: actin sliding velocity was reduced by ∼ 80% compared to WT (*V*_WT_ = 979.6 ± 61 nm/s, *V*_G768R_ = 194 ± 2 nm/s, at 23 °C) (Fig. 1D, Fig. S2). Considering that HCM patients are heterozygous with a mix of mutated and non-mutated proteins in their ventricular muscle, we then repeated the motility assay by titrating increasing proportions of G768R into a background of WT myosin. We found that the reduction in sliding velocity was non-linear and disproportionately large (Fig. 1E). Specifically, the half-max effect (velocity of ∼550 nm/s, halfway between the max of ∼900 nm/s and min of ∼200 nm/s) was observed with only 25% mutant, and velocity had only marginal decreases beyond 40% mutant. These results suggest a dominant effect of the mutation over WT.

Under the detachment-limited conditions of our motility assay, velocity *V* can be approximated as the step size *d* divided by the actin-bound time *t*_b_, which is inversely proportional to the actin-detachment rate *k*_det_: *V* ∼ *d*/*t*_b_ = *d*\**k*_det_. To understand the contributions of these parameters on the mutant’s slow velocity, we next measured the step size and detachment kinetics of single myosin molecules using optical tweezers.

### G768R severely impairs load-sensitive actin detachment kinetics and step size of single myosin molecules

Force-dependent actin-detachment kinetics *k*_det_ of individual sS1 molecules were measured by optical tweezers using the Harmonic Force Spectroscopy (HFS) method and fit to a harmonic force-corrected Arrhenius equation (Eq. 1) (*50*) to extract the zero-load detachment rate (*k*₀) and force sensitivity (*δ*):

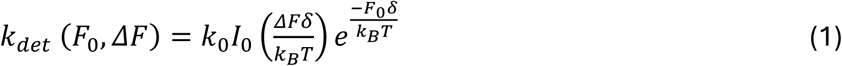

where *F*_0_ is the mean load force, *k*_B_ is the Boltzmann’s constant, T is temperature, and *I*_0_ is the zeroth-order modified Bessel function of the first kind used to correct for the sinusoidal force with amplitude *ΔF*.

The G768R mutation dramatically reduced myosin’s actin detachment rate across all assistive and resistive load conditions (Fig. 2A). The zero-load detachment rate (*k*_0_) was decreased >90%, from 111 ± 10 s−¹ for WT sS1 to 6.7 ± 0.7 s−¹ for G768R (mean ± SEM) (Fig. 2B), indicating a substantially prolonged actin-bound state (WT *t*_b_ ∼10 ms, G768R *t*_b_ ∼150 ms on average). The force sensitivity (*δ*) was also slightly reduced, from 0.88 ± 0.08 nm for WT to 0.71 ± 0.06 nm for G768R (p < 0.001) (Fig. 2C), suggesting altered mechanosensitivity. Finally, the step size of the power stroke, determined from the same HFS data for each molecule (*17, 34, 51*), was decreased by >50%, from 4.1 ± 0.2 nm (WT) to 1.4 ± 0.1 nm (G768R) (p ≤ 0.0001; Fig. 2DE), further indicating impaired mechanical transduction. Consistent with the model *V* ∼ *d*/*t*_b_ = *d*\**k*_det_, G768R’s much reduced actin-detachment rate and step size observed at the single-molecule level explain its slow motility velocity observed at the ensemble level.

**Figure 2.**
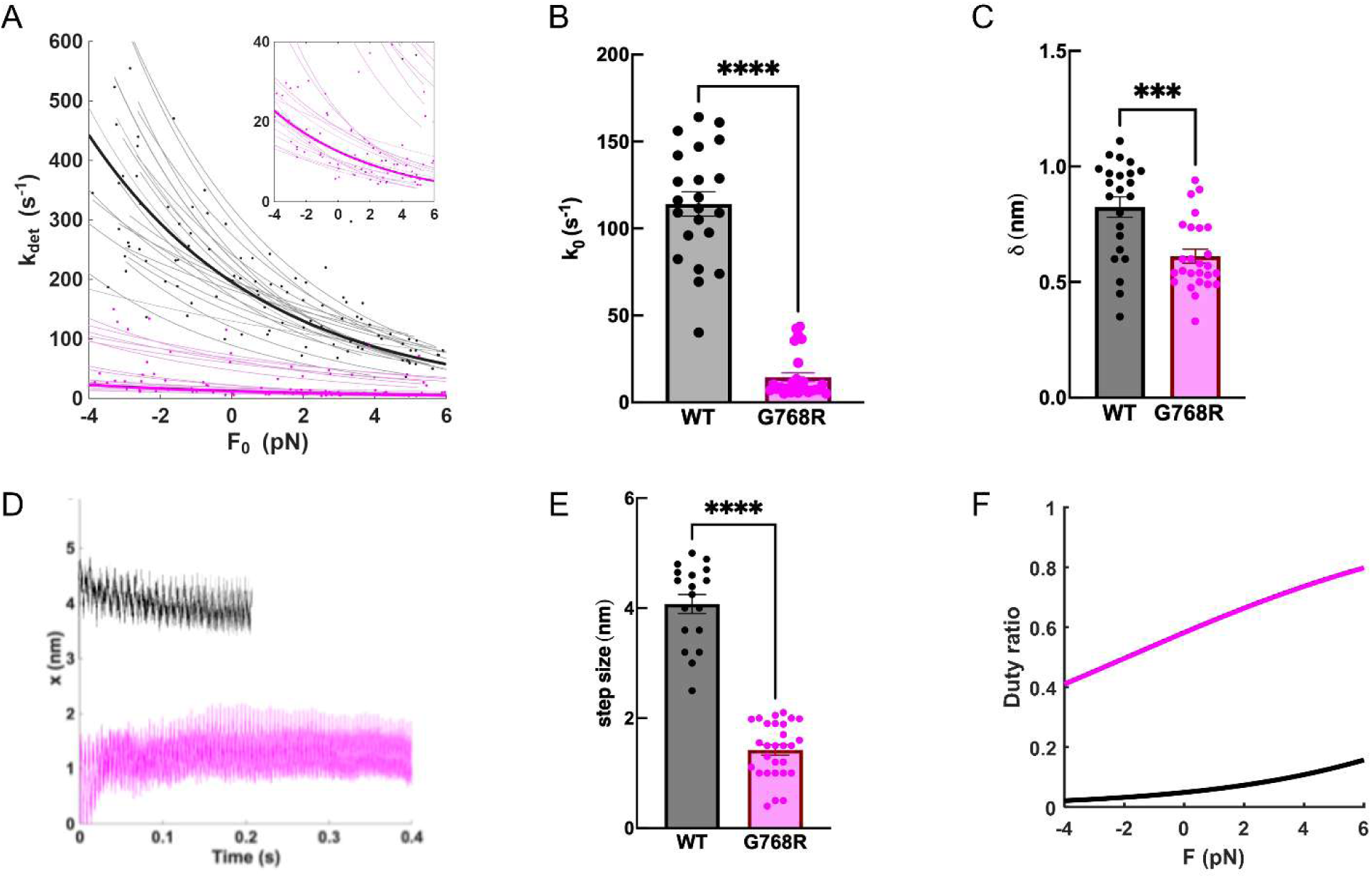
G768R severely slows actin-detachment kinetics of single myosin molecules to make cardiac myosin a high duty ratio motor. **(A)** Load-sensitive actin-detachment rates *k*_det_(F_0_) of WT (black) and G768R (pink) sS1 molecules measured by the harmonic force spectroscopy (HFS) technique in a dual-beam optical trap. Positive forces represent resistive loads, and negative forces represent assistive loads. Each thin line is a fit of Eqn. 1 to data from one molecule, each with a few hundred binding events. Individual data points represent the detachment rates obtained by a MLE (maximum likelihood estimation) of a molecule’s events binned by force. **(B)** and **(C).** The fitted parameters *k*_0_ (detachment rate at zero load) and *δ* (load sensitivity) of each molecule corresponding to thin lines in A. Mean and SEM: WT *k*_0_ = 111 ± 10 s^−1^, *δ* = 0.88 ± 0.08 nm; G768R *k*_0_ = 6.7 ± 0.7 s^−1^, *δ* = 0.71 ± 0.06 nm. **(D)** Averaged, sine-subtracted traces across all binding events of one example molecule each of WT and G768R, from which step size is determined. **(E)** Step sizes of all WT and G768R molecules. Each point represents one molecule. Error bars represent SEM. **** indicates p≤0.0001. **(F)** Calculated duty ratio across different load forces based on measured ATPase and detachment rates (Eqn. 2).

### G768R makes cardiac myosin a high duty ratio motor

From the measured actin detachment rate *k*_det_ and ATPase *k*_cat_, we can next calculate the duty ratio, the fraction of the total cycle time that myosin spends in the actin-bound force-producing state. Since *k*_det_ depends on load force, duty ratio is also a function of load force *F*:

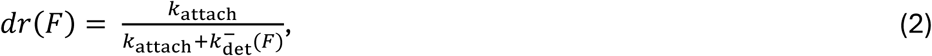

where 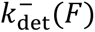 is the detachment rate calculated without harmonic force correction since we generalize to the non-oscillatory case:

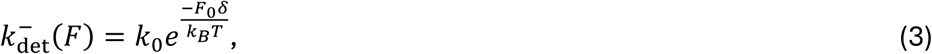

and the attachment rate *k*_attach_ is calculated at saturating actin concentrations and assumed to be independent of force:

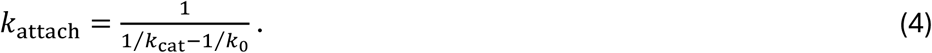

The calculated duty ratios of both WT and G768R sS1 increase as load force increases (Fig. 2F) since the *k*_det_ of both are force dependent: myosin remains bound to actin longer under increasing resistive loads. WT has a duty ratio of ∼0.05 at zero load, as expected of cardiac myosin and consistent with previous studies (*15–17, 52, 53*). In contrast, G768R’s duty ratio is ∼0.6 at zero load, an over 10-fold increase from WT. This huge increase can be attributed to G768R’s substantially slowed *k*_det_ while *k*_cat_ was only slightly decreased, resulting in myosin heads that stay bound to actin for over half its catalytic cycle. Such a magnitude of increase in duty ratio has not, to our knowledge, been previously observed in muscle myosins. This exceptionally high value warrants emphasis, as a muscle myosin that remains actin-bound for more than half of its cycle is bound to exert substantial effects on contractile and relaxation functions and thus carry significant clinical implications.

### G768R disrupts the autoinhibited off-state of myosin

To evaluate the impact of G768R in the context of a two-headed myosin construct, we analyzed heavy meromyosin (HMM) constructs containing 2 or 25 heptad repeats of the tail. The 2-heptad (2-hep) HMM construct does not have a sufficiently long tail to allow the blocked head of the autoinhibited conformation, known as the interacting-heads motif (IHM), to fold back onto its own proximal coiled-coil tail. The 25-hep construct contains a longer coiled-coil tail and *i9* capable of adopting the folded-back IHM conformation (*54, 55*).

The 25-hep WT myosin construct displayed a ∼45% reduction in actin-activated ATPase activity compared to the 2-hep construct (25-hep *k*_cat_ = 3.0 ± 0.3 s^−1^, 2-hep *k*_cat_ = 5.7 ± 0.4 s^−1^) (Fig. 3A, S1, Table S1), consistent with our previously published studies and suggesting that ∼half of WT heads are unavailable to interact with actin (*16, 31, 32, 34, 35*). However, for the G768R myosin, the difference in ATPase activity between the 2-hep and 25-hep constructs was markedly attenuated with only ∼8% decrease (25-hep *k*_cat_ = 3.5 ± 0.4 s^−1^, 2-hep *k*_cat_ = 3.8 ± 0.4 s^−1^) (Fig. 3B, S1, Table S1), indicating that the mutation may open up sequestered heads to be free to interact with actin.

**Figure 3:**
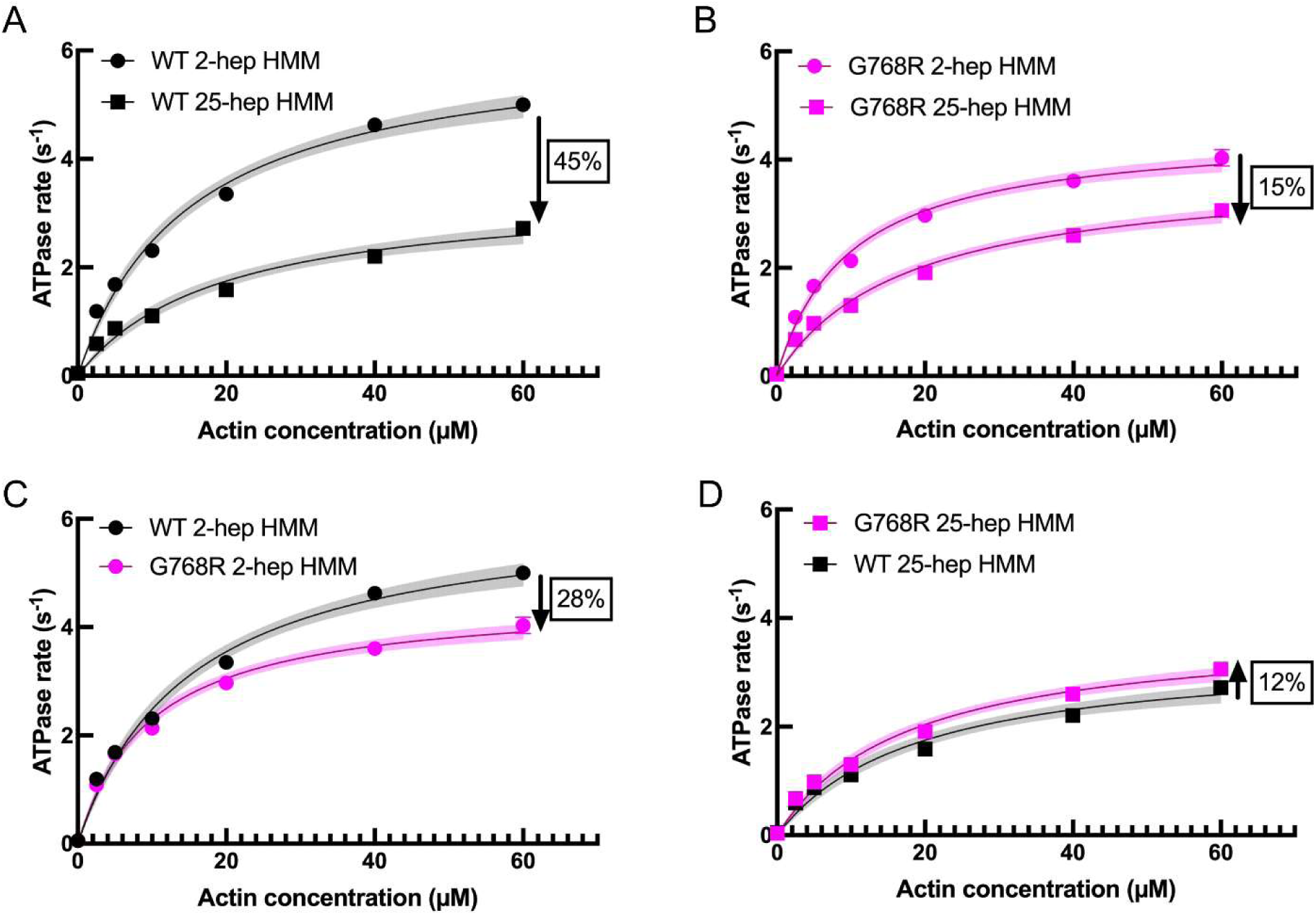
G768R increases the actin-activated ATPase rate of 25-hep HMM β-cardiac myosin by releasing more heads to interact with actin. **(A,B)** Actin-activated ATPase of 2-hep and 25-hep HMM constructs of WT (A) and G768R (B). **(C,D)** Data replotted from (A,B) comparing G768R and WT for 2-hep (C) or 25-hep (D). Each data point represents the average of three technical replicates of one biological replicate and error bars represent SDs. See Table S1 and Fig. S1 for fitted *k*_cat_ and *K*_m_ values, data from other biological replicates, and statistical analyses. Curves are fitted to Michaelis-Menten kinetics, and shaded areas depict the 95% CI of the fits. Arrows with percentages represent changes relative to 2-hep (A,B) or WT (C,D).

Interestingly, the 2-hep G768R construct exhibited a reduced ATPase rate relative to WT (Fig. 3C) (similar to the results obtained with sS1, see Fig. 1C), whereas the 25-hep G768R construct showed a slight increase in ATPase activity (Fig. 3D). This pattern suggests that the enhanced availability of previously sequestered heads in the mutant 25-hep construct compensates for the reduced catalytic activity of the mutant and may be a reason this HCM mutation results in hypercontractility of the heart.

### Simulations reveal structural mechanisms for inhibition of power stroke and ADP release by G768R

We performed MD simulations of WT and G768R sS1 starting from homology models built on a pre-powerstroke (PPS) (PDB: 8QYU) and a post-rigor (PR) (6FSA) β-cardiac myosin crystal structure. Eight independent 1-2 µs simulations were performed for each of the four initial models: WT PPS, WT PR, G768R PPS, and G768R PR (Fig. S3). The models are comprised of the heavy chain (residues 2-810) and the essential light chain (ELC) (Fig. 4A).

**Figure 4.**
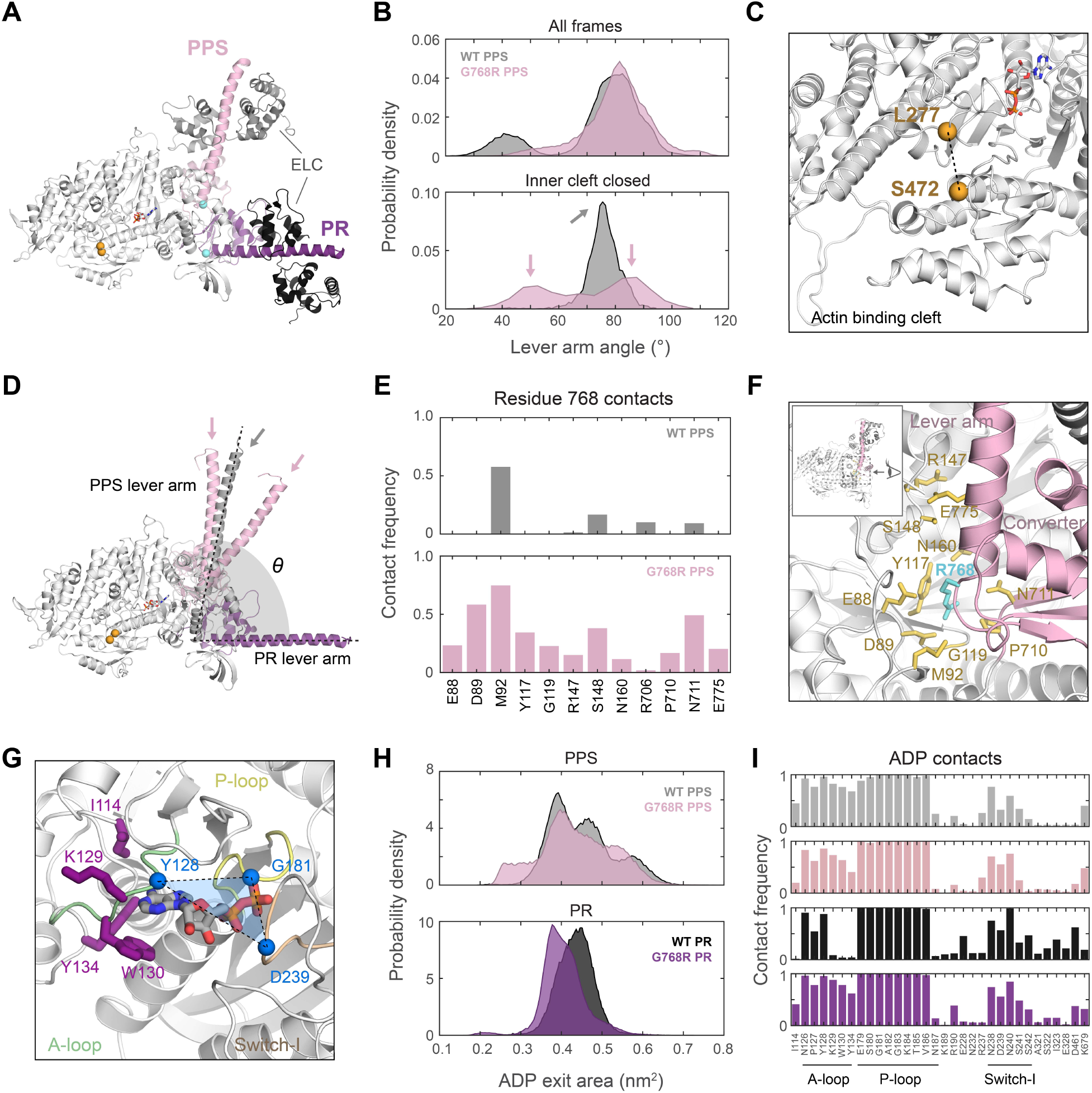
The G768R mutation alters lever arm priming and reduces ADP pocket opening in MD simulations of β-cardiac myosin. **(A)** Initial structural models of the human WT and mutant β-cardiac myosin were built from a pre-powerstroke (PPS) (PDB: 8QYU) and a post-rigor (PR) (6FSA) WT structure. The models are comprised of the heavy chain head (residues 2-810) and the essential light chain (ELC). The motor domain is colored white; the converter domain and lever arm are colored pink in PPS and purple in PR models. Residue 768 is shown as a cyan sphere. **(B)** Probability distributions of the lever arm angle (as defined in **(D)**) in all PPS simulations (top) and conditioned on inner cleft closed (≤ 1.1 nm) (bottom). Among all frames, G768R has an overall population shift towards larger angles compared to WT. Among cleft-closed states, the mutant also exhibits a smaller angle population suggesting loss in lever arm priming. **(C)** Opening and closing of the actin binding cleft is observed in the simulations of models in the PPS conformation. The inner cleft distance, between the α-carbons of L277 and S472, is used as a metric of cleft closure ( Liu et al., 2024). **(D)** The lever arm angle was defined as the angle between the lever arm in simulations and the PR crystal structure (see Methods). The lever arms in gray and pink represent three simulated instances corresponding to the peaks of the angle distributions of WT and G768R models, respectively (arrows in **(B)**). **(E)** Contacts between residue 768 and other residues are shown only if their frequency is higher than 10% in the PPS simulations. The arginine mutation gains electrostatic and polar interactions with nearby residues in the N-terminus and converter domain. **(F)** Contacts gained by R768 are shown in yellow licorice and are located to stabilize conformations with larger lever arm angle in the PPS state. **(G)** The ADP pocket exit area is defined as a triangle formed by the α-carbons of Y128, G181, and D239 (blue). **(H)** Probability distributions of the ADP exit area. WT PPS simulations sample both pocket-closed (∼0.4 nm^2^) and pocket-open (∼0.6 nm^2^) states. G768R PPS simulations have a shift away from the pocket-open peak and sample smaller exit area states more often. **(I)** Contact between ADP and myosin residues are shown only if their frequency is higher than 10%. Residues I114, K129, W130, and Y134 (purple licorice in **(G)**) show the most significant gain of contact in G768R PR models compared to WT PR.

In simulations started from the PPS models, the lever arm remained for the majority of the time in an up position with angle ∼80° relative to the lever arm of the post-rigor crystal structure, representing the fully primed state (Fig. 4B top). While WT PPS simulations also explored lower lever arm angles (∼45°), towards the post-stroke state, the lever arm in G768R PPS simulations lacked a prominent distribution peak at these smaller angles and instead had an overall shift towards larger angles (Fig. 4B, S4). The inability to explore lever arm down states in simulations suggests that the G768R mutation may keep myosin in a high-angle primed state and prevent the execution of the power stroke, thus explaining our experimental finding of a smaller step size.

To further explore this proposed mechanism, we analyzed the position of the lever arm in the context of actin binding cleft behavior. An inner cleft distance, defined between the α-carbons of L277 and S472 (Fig. 4C), of ≤ 1.1 nm was used as a metric of cleft closure (*17, 56*). Opening and closing of the actin binding cleft was observed in simulations. Cleft-closed states occupied a very small fraction (∼2-7%) of all PPS simulation frames, as expected since these models did not involve actin. Nevertheless, since cleft closure is required for strong actin binding and a productive power stroke, an analysis of the behavior of the lever arm when the cleft is closed can provide clues to myosin’s action if actin were present. We found that when the cleft was closed, the lever arm angle distribution had a single peak at ∼75° in WT PPS simulations (Fig. 4B bottom, 4D), in close agreement with previous simulation results (*17*). The lower angle peak of ∼45° that was present when the actin cleft was open (Fig. 4B top) was now absent. This is consistent with our expectations because the lever arm should be locked into the fully primed position and ready to stroke when the cleft is closed, mimicking the actin-bound pre-stroke state.

In contrast, we found that G768R occupied a lower angle state of ∼50° in almost half of the cleft-closed PPS simulation frames (Fig. 4B bottom, 4D). This significant loss in lever arm priming is expected to result in a less productive lever arm swing consistent with G768R’s experimentally observed reduced step size. Interestingly, the remaining fully-primed high-angle population was shifted higher than WT to ∼85° (Fig. 4B bottom, 4D). However, we found that the lever angle of cleft-closed PR simulations was also larger (towards pre-stroke) for G768R than WT (Fig. S5). This finding, along with the absence of the lever-down peak in the total simulation distribution that was present in WT (Fig. 4B top), suggests that the mutation may hinder a full power stroke, resulting in a smaller step size despite having a population with larger primed angle.

To examine how a glycine to arginine substitution at position 768 may lead to higher lever arm angles, we examined the frequencies of contacts between residue 768 and other myosin residues across all independent simulations carried out for the models (Fig. S6). Two residues were defined to be in contact when the distance between their non-hydrogen atoms was 0.5 nm or less. In comparison to G768, R768 in PPS simulations gained 10 additional contacts with surrounding residues (Fig. 4E). Seven of those are in the N terminal domain (E88, D89, Y117, G119, R147, S148, and N160) and two are at the start of the converter domain (P710, N711) (Fig. 4F). Due to the location of these residues, the increased electrostatic and polar interactions between them and R768 are expected to pull the lever arm towards a more primed position, stabilizing conformations with larger lever arm angles as observed in G768R PPS simulations (Fig. 4B, D).

Finally, we examined the behavior of the nucleotide binding pocket in both PPS and PR simulations and found that it adapted conformations that are predicted to hinder ADP release. The ADP pocket exit area is defined as a triangle between the A-loop, P-loop, and Switch-I (Fig. 4G) (*17*). WT PPS simulations sampled both pocket-closed (∼0.4 nm^2^) and pocket-open (∼0.6 nm^2^) states (Fig. 4H top). In contrast, G768R PPS simulations had a shift away from the pocket-open peak and sampled smaller exit area states more often. We observed a similar trend in PR simulations: the distribution of the ADP exit area is much smaller in G768R simulations than in WT (Fig. 4H bottom). As the release of ADP is a prerequisite for ATP binding and subsequent actin detachment, this finding may explain G768R’s much reduced actin-detachment rate observed in our single-molecule trapping experiments.

To further explore this effect, we examined frequences of contacts between ADP and residues of the nucleotide binding pocket. Although there were no large differences between WT and G768R in PPS simulations, G768R PR simulations showed significant gain of contact by residues I114, K129, W130, and Y134 in or near the A-loop compared to WT (Fig. 4G, I; Fig. S7). We note that neither of the two starting models (PPS and PR) are the post-stroke, actin-bound, pre-ADP release state (as no crystal structures exist for this state) which would have been the most informative conformation for studying the release of ADP. Nevertheless, our findings of smaller ADP exit area and gained ADP contacts in two different models suggest that the G768R mutation may induce allosteric changes to the nucleotide binding pocket’s local structure and nucleotide-myosin interactions that slow the release of ADP and thus actin detachment as observed experimentally.

## Discussion

The findings presented herein elucidate the profound mechanistic disruptions imparted by the G768R mutation in human β-cardiac myosin, underscoring its pathological potential in HCM. Situated at the distal converter domain immediately proximal to the lever arm—a region crucial for coupling chemical and mechanical transitions—the G768R substitution exerts consequential allosteric changes in myosin kinetics, mechanosensitivity, and regulatory state transitions.

Steady state actin-activated ATPase measurements (Fig. 3) on HMM constructs containing the regulatory tail suggest that the G768R mutation reduces the population of myosin in the sequestered autoinhibited “OFF” state. Thus, G768R lends yet another example in support of the unifying hypothesis proposed by Spudich in 2015 that most HCM myosin mutations cause an increase in the number of heads available for interaction with actin (*57*). The magnitude of G768R’s effect on ATPase rate (15% lower for 25-hep vs 2-hep, vs. 45% lower in WT) is similar to or modest in comparison to changes previously observed for other mutations, some of which had 25-hep ATPase as fast as or faster than 2-hep ATPase (*14, 16, 31, 32, 34, 35*).

In contrast, G768R’s effect on motor actin-detachment kinetics was much more dramatic. The precipitous drop in the detachment rate in the mutant relative to WT myosin by over an order of magnitude (from *k*_0_ ∼100 to 6.7 s^−1^) reflects a substantial prolongation of the actin-bound state (from *t*_b_ ∼ 1/*k*_0_ ∼10 to 150 ms), as uncovered by single molecule optical trapping experiments (Fig. 2). This magnitude of change has been observed previously only when myosin is treated with small molecule drugs like omecamtiv mecarbil (OM) (*15, 58*) or in severe fetal skeletal myosin mutations (*17*), but not in cardiac myosin mutations.

The marked increase in actin-bound time coupled with only a modest increase in the total actin-activated ATPase cycle time (WT *k*_cat_ = 5.5 s^−1^, *t*_cycle_ ∼ 180 ms vs. G768R *k*_cat_ = 4.0 s^−1^, *t*_cycle_ ∼ 250 ms, Fig. 1C) transforms cardiac myosin into a high duty ratio (>0.6) motor. This increase in duty ratio is significantly greater than that observed for any previously published mutations or myosin modulatory drugs. Published single-molecule and biochemical studies report duty ratios of ∼0.05 for WT cardiac myosin and most myosin mutations, with only a few select mutations and activators such as OM increasing the duty ratio to at most ∼0.2 (*15–17, 59–62*). Thus, G768R’s markedly higher duty ratio of ∼0.6 underscores the unusually long actin attachment of this mutant. In the heart, such prolonged force-generating states would be expected to cause persistent thin filament activation, delayed relaxation, and thus perturbed diastolic function, all hallmarks of HCM pathology.

These molecular changes at the single molecule level manifest clearly at the ensemble level. Consistent with the simple model *V* ∼ *d*/*t*_b_ = *d*\**k*_det_, G768R’s much reduced actin-detachment rate and step size explain its striking ∼80% reduction in motility velocity (Fig. 1D). This relationship has been previously confirmed for many other mutations (see Liu PNAS 2024 (*17*) for a compilation). Furthermore, the disproportionally severe effect of the mutation in the mixed G768R/WT motility experiments (Fig. 1E), mimicking the heterozygous genotype in HCM patients, can be attributed to the dramatically slowed actin-detachment kinetics of the mutant. In these experiments, the G768R subpopulation of myosin heads that are bound to actin for a long time among a sea of WT heads exert a disproportionally large effect to slow down velocity. This is similar to the common effect observed in motility assays in which a very small proportion of dead heads cause a huge reduction in velocity and increase in stuck actin filaments (see discussions in Schroer PNAS 2021 (*34*) and Liu PNAS 2024 (*17*)). Thus, G768R likely exerts a dominant-negative effect beyond its stoichiometric representation.

The effects of the G768R mutation on β-cardiac myosin function share similarities with the Freeman-Sheldon Syndrome (FSS, a congenital disorder with severe muscle contractures) R672C mutation in embryonic skeletal myosin (MYH3 gene) (*63*), which we had previously studied using similar experimental and computational analyses (*17*). Both mutations had slowed actin-detachment rate as measured by single molecule optical trapping (Fig. 2), and MD simulations suggested that a bias towards a smaller or closed ADP exit area may be responsible (Fig. 4H). Both mutations were also measured by single molecule experiments to have reduced step sizes (Fig.2), and simulations suggest that a significant population with smaller primed lever arm angle among actin binding cleft-closed states can explain this reduction (Fig. 4B bottom). While R672C also exhibits a population with lever angles only marginally higher than WT, the higher angle population in G768R is more pronounced (Fig. 4B bottom), possibly due to electrostatic interactions gained by the arginine mutation with surrounding N-terminal and converter domain residues (Fig. 4EF). However, G768R’s power stroke may be inhibited or reduced, as suggested by the lack of sampling low lever angle conformations among all PPS simulation frames compared to WT (Fig. 4B top) and the observation of higher lever angles among cleft-closed states in PR simulations (Fig. S5). Taken together, simulations suggest two possible mechanisms to explain G768R’s experimentally measured smaller step size: the mutation reduces PPS lever arm priming but also stabilizes a high-angle lever arm conformation that is hindered from fully stroking.

The effects of G768R, in both experimental and simulation results, are also reminiscent of the investigational myosin modulatory drug omecamtiv mecarbil (OM). Both G768R and OM have a small or close to zero measured step size (*51, 58*), a larger primed lever arm angle in PPS simulations (*64*) and in the crystal structure of OM-bound WT myosin (*65, 66*), and prolonged actin bound time (*15, 58*). Of the contacts gained by the arginine mutation at position 768 according to MD simulations, four (P710, N711, R147, and N160) are key residues involved in binding OM (*65, 66*). This suggests that the G768R mutation may act through a mechanism similar to OM. Importantly, three of these contacts (P710, R147, and N160) gained by 768R are *not* residues involved in binding the FDA-approved myosin inhibitor Mavacamten (Mava), despite both OM and Mava sharing the same binding pocket (*65*). Unlike OM, Mava does not change myosin’s step size or actin-bound time (*67, 68*), and it also does not stabilize the lever arm in the highly primed position to the same extent (*65*). This contrast between Mava and OM/G768R is a testament to myosin’s highly allosteric and specific nature: only perturbations to specific residue interactions, shared by OM and 768R but not Mava, in a common binding pocket can stabilize a primed lever arm, inhibit the power stroke, and slow down actin detachment.

Comparisons to both R672C and OM provide insight into the pathophysiology of G768R in the heart. Specifically, both provide evidence to the hypothesis that a myosin with a significantly prolonged actin-bound time in vitro impairs the relaxation of muscle in patients. The FSS mutation R672C doubles the bound time of embryonic skeletal myosin (*17*), and muscle cells and myofibrils isolated from R672C FSS patients had profoundly slowed relaxation (*69*). OM increases the bound time of β-cardiac myosin by up to 10x in a dose dependent manner (*15*), and patients given high doses experienced ischemic effects from excessively prolonged systole and impaired relaxation (*70*). By the same token, G768R’s profound prolongation of the bound time in β-cardiac myosin by >10x is expected to slow relaxation of the heart and thus cause diastolic dysfunction, a hallmark of HCM. The severity of this mutation’s effect on cardiac function may explain its very low occurrence in the population and its early-onset, severe manifestation (*48, 49*). Furthermore, HCM patients, who are all heterozygous, carrying this mutation may tolerate only low expression of the mutant protein since it exerts a disproportionally large effect (Figure 1E).

In summary, the G768R mutation exerts a dual-axis disruption: it compromises detachment kinetics and mechanics of the motor, while simultaneously undermining the autoinhibitory regulatory architecture that governs cardiac efficiency. The resulting high duty ratio and loss of autoinhibition together promote a more functionally engaged and energetically costly myosin population. Impaired relaxation of the heart is expected as a consequence, a hallmark of HCM disease. Such mechanistic insight provides a framework for understanding how seemingly subtle point mutations precipitate complex clinical phenotypes. It also offers fertile ground for therapeutic intervention—whether through small molecules that restore proper detachment kinetics or ones that re-stabilize the autoinhibited state.

## Methods

### Experimental methods

Recombinant human β-cardiac myosin constructs—including short subfragment 1 (sS1), short-tailed (2-hep HMM), and long-tailed (25-hep HMM) 2-headed forms—were expressed in C2C12 mouse myoblasts and purified as previously described (*16, 34*).

Load-dependent detachment rates for WT and G768R sS1 were measured using harmonic force spectroscopy (HFS) in a dual-beam optical trap, following established protocols (*15, 50*). Experiments were done at room temperature (23 °C). Myosin step sizes were derived from the same HFS datasets using an adapted ensemble averaging approach suitable for oscillatory data (*17, 51, 71*). For each molecule, binding event traces were aligned at binding onset, extended to match the longest event, averaged, and then de-oscillated by subtracting a fitted sine wave. Step size was defined as the displacement between the initial and final positions of these processed traces.

In vitro motility assays of WT and G768R sS1-AC were performed as previously described (*72, 73*) with minor modifications. Mixed in vitro assays were performed as described previously (*72*). All experiments were done at room temperature, with the temperature of the imaging microscope maintained at 23 ± 1 °C. Velocities reported in main figures represent MVEL20 from FAST analysis of motility movies, with parameters: window size n=5, path length p=10, percent tolerance pt=20, and minimum velocity threshold for stuck filament classification minv=80 nm/s. See Aksel Cell Rep 2015 (*72*) for the original development of the FAST analysis program and the Supplement in Liu PNAS 2024 (*17*) for detailed explanations of different velocity parameters.

NADH-coupled ATPase assays were used to assess enzymatic activity in WT and G768R 2-hep and 25-hep constructs, following established methods (*16, 74*). All experiments were done at room temperature (23 °C). Paired (one-sample) t-test is used to calculate p values for statistical significance between WT and G768R (Figure S1, Table S1).

### MD simulation methods

The β-cardiac myosin structural models were built with Modeller 10.5 (*75*) templated on the crystal structures of a bovine (97.6% sequence identity to the human MYH7 protein) pre-powerstroke (PDB: 8QYU) (*65*) and a post-rigor (6FSA) (*76*) state. The resulting models included residues 2-810 of myosin heavy chain, residues 39-195 of the light chain, ADP, and Mg^2+^ for the PR and PPS models, and inorganic phosphate for PPS models. The models were solvated with 0.1M NaCl with CHARMM GUI’s Solution Builder (*77*). The solution box extended 1 nm beyond the protein in every dimension, for a total of ∼ 370K – 418K atoms within a ∼15.5 nm x 15.5 nm x 15.5 nm box. After a 5000-step energy minimization with steepest descent algorithm, a three-step equilibration was carried out in the NVT ensemble, with temperature kept at 310 K with a v-rescale thermostat (*78*). Production runs were performed with all the positional restraints removed, and temperature and pressure were kept at 310 K and 1 bar using v-rescale thermostat and c-rescale barostat (*79*). CHARMM36m force field (*80*) with TIP3P water model (*81*) were used. For ADP and Pi, CHARMM General Force Field parameters were taken from CHARMM GUI’s library. The simulations were carried out with GROMACS 2023.4 (*82*).

Using this protocol, we performed 8 independent simulations (replicas) each for WT PPS, WT PR, G768R PPS, and G768R PR (32 simulations total). The aggregated simulation times were 11.6 µs for WT PPS, 11.2 µs for WT PR, 11.2 µs for G678R PPS, and 10.9 µs for G768R PR. The first 120-250 ns of each replica simulation was discarded for further equilibration of the simulation systems. The analyzed part of the trajectories had an aggregated time of 9.7 µs for WT PPS, 9.4 µs for G768R PPS, 9.4 µs for WT PR, and 9.2 µs for G768R PR.

Analyses of the lever arm angle and ADP exit area followed a previous work (*17*). Contact frequencies were determined from distances between non-hydrogen atoms of ADP (or residue 768) with myosin residues: if the distance was 0.5 nm or less, we defined those residues as in contact (Fig. S6 and S7). Frequences of >0.1 are shown in the main figures.

Additional computational details are in supplementary methods.

## Author contributions

D.P. performed experiments and analyzed data. T.N.O performed simulations and analyzed the simulation data. D.P. and C.L. wrote the paper. M.R., J.A.S., C.L., and K.M.R. supervised the work. All authors contributed intellectually to discussions of the results and reviewed and edited the paper.

## Acknowledgements

D.P., K.M.R., and J.A.S. were funded by NIH R01 GM033289 and RM1 GM131981 to J.A.S. D.P was funded by AHA Postdoctoral Award #831317. T.N.O. and C.L. were funded by Lawrence Livermore National Laboratory LDRD 25-LW-138 awarded to C.L. T.N.O. thanks the Livermore Institutional Grand Challenge for computing time. The microscope was funded by NIH Grant S10RR026775 to J.A.S. M.R. was funded by NIH RM1 GM131981, R01HL179584, and P30AR074990. M.C.C. was funded by NIH K99 HL173646. Part of this work was performed under the auspices of the U.S. Department of Energy by Lawrence Livermore National Laboratory under Contract DE-AC52-07NA27344, release number LLNL-JRNL-RR0153233.

