## Supporting Information for "The hypertrophic cardiomyopathy mutation G768R makes cardiac myosin a high duty ratio motor"

**Supplementary methods**

**Protein expression and purification**

Recombinant human β-cardiac myosin constructs, including sS1, 2-hep HMM (short-tailed), and 25-hep HMM (long-tailed), were purified using previously described methods (*1, 2*) with minor modifications. Human cardiac myosin heavy chain (MYH7) was co-expressed with human essential light chain (ELC; MYL3) containing an N-terminal FLAG tag followed by a tobacco etch virus (TEV) protease cleavage site. Expression was performed in differentiated mouse myoblast C2C12 cells (ATCC) using adenoviral vectors generated in HEK293T cells (ATCC) via the AdEasy Vector System (Qbiogene Inc., Carlsbad, CA, USA). The sS1 construct contained a C-terminal enhanced green fluorescent protein (eGFP) tag, whereas the 2-hep and 25-hep constructs contained both eGFP and a PDZ C-peptide at their C-termini.

C2C12 cells were infected with adenoviral constructs 48 hours post-differentiation and harvested 4 days post-infection. Cells were lysed in buffer containing 50 mM NaCl, 20 mM $MgCl_2$, 20 mM imidazole (pH 7.5), 1 mM EDTA, 1 mM EGTA, 1 mM DTT, 3 mM ATP, 1 mM phenylmethylsulfonyl fluoride (PMSF), 5% sucrose, and cOmplete protease inhibitor cocktail (Roche). This lysis buffer formulation differed from our previously published methods in that salt concentration and sucrose were reduced while $MgCl_2$ concentration was increased to promote native mouse myosin filament formation, thereby reducing contamination from murine myosin. Following lysis, cells were immediately flash-frozen in liquid nitrogen. Cell pellets were stored at –80°C for up to 6 months prior to purification.

For purification, frozen pellets were thawed at room temperature and subjected to 50 strokes in a Dounce homogenizer on ice. Lysates were clarified by ultracentrifugation at 30,000 rpm in a Ti-60 fixed-angle rotor for 30 minutes at 4°C. The supernatant was incubated with anti-FLAG affinity resin for 1–2 hours at 4°C. The resin was then washed with buffer containing 150 mM NaCl, 5 mM $MgCl_2$, 20 mM imidazole (pH 7.5), 1 mM EDTA, 1 mM EGTA, 1 mM DTT, 3 mM ATP, 1 mM PMSF, 10% sucrose, and cOmplete protease inhibitor cocktail (Roche).

For the 2-hep HMM and 25-hep HMM constructs, endogenous mouse regulatory light chain (RLC) was depleted by incubating the resin with buffer containing 20 mM Tris (pH 7.5), 200 mM KCl, 5 mM CDTA (pH 8.0), and 0.5% Triton X-100 for 1 hour at 4°C. Recombinant human RLC, purified from Escherichia coli as previously described (*1*), was then added to the resin in wash buffer and incubated for at least 2.5 hours at 4°C. The ELC-myosin complex was subsequently released from the resin by overnight incubation with TEV protease at 4°C.

The cleaved protein was further purified by anion-exchange chromatography using a HiTrap Q HP column on a fast protein liquid chromatography (FPLC) system with a linear gradient of 0–600 mM NaCl over 20 column volumes in buffer containing 10 mM imidazole (pH 7.5), 4 mM $MgCl_2$, 10% sucrose, 1 mM DTT, and 2 mM ATP. Fractions containing pure protein, as assessed by Coomassie staining of 10% SDS-PAGE gels, were pooled and concentrated to 5–50 μM using Amicon Ultra 0.5 mL centrifugal filters with a 50 or 100 kDa molecular weight cutoff, which also facilitated removal of unbound ELC or RLC. Purified myosin was either used immediately for ATPase assays, buffer-exchanged for single-turnover experiments, or flash-frozen in liquid nitrogen and stored for subsequent in vitro motility or optical trapping experiments.

**Actin-activated ATPase assay**

Actin-activated ATPase activity was measured using an NADH-coupled enzymatic assay based on established protocols (*3*). Actin was purified, polymerized, and dialyzed four times against assay buffer containing 5 mM KCl, 10 mM imidazole (pH 7.5), 3 mM $MgCl_2$, and 1 mM DTT. The dialyzed actin was then mixed with bacterially expressed and purified gelsolin at a 1:100 molar ratio, thoroughly combined, and incubated on ice for at least 30 minutes.

Assays were performed in clear 96-well plates with a final reaction volume of 100 μL per well. The actin-gelsolin mixture was diluted in assay buffer to generate final actin concentrations of 0–80 μM, and myosin was added to a final concentration of 25 nM. For basal ATPase activity measurements, myosin alone was added at 75–125 nM in the absence of actin. Plates were pre-incubated at room temperature for 10 minutes with continuous shaking. Reactions were initiated by adding 20 μL of 5× coupling solution containing 100 U/mL lactate dehydrogenase (Sigma-Aldrich #L1254, St. Louis, MO, USA), 500 U/mL pyruvate kinase (Lee Biosolutions #500–20, Maryland Heights, MO, USA), 2.5 mM phosphoenolpyruvate (Sigma #P0564), 10 mM ATP, and 2 mM NADH (Sigma #N8129). Following a further 2–5 minute incubation at room temperature with shaking, absorbance at 340 nm was measured every 15–30 seconds for 15–25 minutes. ADP concentration was calculated using a standard curve generated with ADP concentrations of 0–300 μM.

ATPase rates at each actin concentration were determined from the linear slope of ADP accumulation over time. Technical triplicates were conducted for each actin concentration on the same plate using identical protein preparations. Rates were normalized to the myosin concentration in each well, plotted as a function of actin concentration, and fitted to Michaelis-Menten kinetics to obtain the reported kcat and Km values. G768R mutant constructs (sS1, 2-hep, and 25-hep) were prepared in parallel with their corresponding wild-type (WT) controls (sS1, 2-hep, and 25-hep, respectively), and technical triplicates were performed across three independent biological replicates, as detailed in Supplementary Table 1 and Supplementary Figure S1. Biological replicates employed proteins derived from different C2C12 cell batches prepared on separate days. Statistical comparisons between WT and mutant ATPase rates were conducted using one sample (paired) Student's t-test. In Supplementary Table 1, each biological replicate was independently fitted to standard Michaelis-Menten kinetics to calculate kcat and Km values; the means and SEMs across the biological replicates are also reported.

**In vitro actin motility assay**

In vitro motility assays were conducted following established protocols (*4*). Multichannel flow chambers were assembled on microscopy slides using double-sided tape and coverslips pre-coated with 0.1% nitrocellulose/0.1% collodion in amyl acetate. Chamber surfaces were functionalized through sequential 2-minute incubations at room temperature as follows: initial incubation with 3 µM SNAP-PDZ in assay buffer (AB; 25 mM imidazole pH 7.5, 25 mM KCl, 4 mM MgCl2, 1 mM EGTA, 10 mM DTT), followed by two blocking steps using AB supplemented with 1 mg/mL BSA (ABBSA). Myosin constructs were diluted to 50–100 nM in ABBSA and incubated for 2 minutes to facilitate surface attachment. Following an ABBSA wash, channels were loaded with the final motility buffer comprising ABBSA, 2 mM ATP, 2.5 nM TMR-phalloidin-labeled actin, and an oxygen-scavenging system (0.4% glucose, 0.11 mg/mL glucose oxidase, 0.018 mg/mL catalase).

Imaging was performed on a Nikon Ti-E inverted microscope equipped with a 100× TIRF objective and Andor iXon+ EMCCD camera. Three fields of view per channel were recorded for 30 s at an acquisition rate of 2 Hz with 300 ms exposure time. Actin filament tracking and velocity quantification were executed using Fast Automated Spud Trekker (FAST) (*4*) with parameters set as follows: window size n = 5, path length p = 10, percent tolerance pt = 20, and minimum velocity threshold for stuck filament classification minv = 80 nm/s. Filtered mean velocity values were used to represent unloaded velocity. Examples of FAST-analyzed data are show in Supplementary Figure S2.

To control for ambient temperature fluctuations (21–23°C), G768R sS1 and wild-type (WT) sS1 were assayed simultaneously on the same slide. Three biological replicates from independent protein preparations were analyzed. Statistical analysis was performed using a paired t-test to compare G768R velocities against their corresponding WT controls.

### MD simulation details

**Model Preparation.** The β-cardiac myosin structural models in the pre-powerstroke (PPS) and post-rigor (PR) conformations were built with Modeller 10.5 (*5*) using the crystal structures taken from the Protein Databank (*6*) with entries 8QYU (*7*) and 6FSA (*8*), respectively. The wild-type (WT) sequences were taken from the UNIPROT (*9*) with entries P12883 for the human myosin 7 and P08590 for the human myosin light chain 3. Bovine and human sequences were aligned with Clustal Omega (version 1.2.4) (*10*). With Modeller, we also introduced the G768R mutation and built the mutant structural models in both PPS and PR conformations. Among the generated 20 models, the best four structures (WT-PPS, WT-PR, G768R-PPS and G768R-PR) were chosen as the initial structures for molecular dynamics simulations. The resulting models included the residues 2-810 of the myosin, residues 39-195 of the light chain, ADP, and $Mg^{2+}$ for those both in the PR and PPS conformations. The PPS structural models also included the inorganic phosphate as it was present in the template structure as a vanadate ion.

**Simulation setup.** The models were solvated with 0.1M NaCl with CHARMM GUI's Solution Builder (*11*); the solution box extended 1 nm beyond the protein in every dimension. The resulting simulation systems had about 370K – 418K atoms within ~15.5 nm x 15.5 nm x 15.5 nm. After a 5000-step energy minimization with steepest descent algorithm, a three-step equilibration was carried out in the NVT ensemble. First, protein backbone and sidechain atoms were equilibrated using positional restraints with force constants of 500 kJ/mol/nm$^2$ and 50 kJ/mol/nm$^2$ for 0.1 ns (time step was set to 1 fs). Then the force constants for the positional restraints were decreased to 200 kJ/mol/nm$^2$ and 20 kJ/mol/nm$^2$ for 0.5 ns with a time step of 2 fs. Finally, only the backbone atoms were restrained with a force constant of 50 kJ/mol/nm$^2$ for 0.5 ns with a time step of 2 fs. During the equilibration steps, the temperature was kept at 310 K with a v-rescale thermostat (*12*). Next, we carried out production runs during which all the positional restraints were removed, and the temperature and pressure were kept at 310 K and 1 bar using v-rescale thermostat and c-rescale barostat (*13*). The rest of the simulation parameters was kept as in the CHARMM GUI's protocol. CHARMM36m force field (*14*) with TIP3P water model (*15*) were used. The simulations were carried out with GROMACS 2023.4 (*16*).

Using this protocol, we performed 8 independent simulations (replicas) each for WT PPS, WT PR, G768R PPS, and G768R PR (32 simulations total). The aggregated simulation times were 11.6 μs for WT PPS, 11.2 μs for WT PR, 11.2 μs for G678R PPS, and 10.9 μs for G768R PR.

**Analyses of simulation trajectories.** To check structural stability, we calculated the root mean square deviation (RMSD) from the initial states using the alpha carbon atoms, the root mean square fluctuations (RMSF) per residue using all non-hydrogen atoms, and the total number of unique hydrogen bonds over the entire production runs (see Supplementary Figure S3). These analyses were done for the myosin and essential light chain separately for each of the eight replicate simulation trajectories. Hydrogen bonds were defined using a distance and angle cut-off values of 0.3 nm and 20° between each donor-acceptor pair.

For the rest of the analyses, the first 120-250 ns of each replica simulation was discarded for further equilibration of the simulation systems. The analyzed part of the trajectories had an aggregated time of 9.7 μs for WT PPS, 9.4 μs for G768R PPS, 9.4 μs for WT PR, and 9.2 μs for G768R PR.

The lever arm angle (Supplementary Figures S4 and S5) was calculated as explained in a previous work (*17*). Briefly, we first aligned the WT PPS and WT PR crystal structures using the alpha carbon atoms of residues 2-760 and defined a plane where their lever arm vectors (formed from the geometric center of alpha carbon atoms of residues 770-776 to the center of residues 800-806) lie. We then calculated the lever arm angle of our models in each snapshot of their simulation trajectories as the angle between their lever arm vector and the lever arm vector of WT PR initial structure on the defined plane. The inner cleft was defined as the distance between the alpha carbon atoms of L277 and S472. The inner cleft was considered closed if this distance was equal or less than 1.1 nm as defined previously.

To calculate contact frequencies, we analyzed the distance between non-hydrogen atoms of ADP (or residue 768) with myosin residues: if the distance was 0.5 nm or less, we defined those residues as in contact (Supplementary Figures S6 and S7).

The ADP exit area was defined as the area of a triangle formed by the alpha carbons of Y128, G181 and D239 for a consistent comparison with a previous work (*17*).

Simulation trajectories were analyzed using MDAnalysis (*18*), VMD (*19*) and GROMACS (*16*) analysis tools. The snapshots were generated with Pymol (The PyMOL Molecular Graphics System, Version 3.0 Schrödinger, LLC).

| | sS1 | | | 2-hep HMM | | | 25-hep HMM | | |
|---|---|---|---|---|---|---|---|---|---|
| | $k_{cat}$ ($s^{-1}$) | $k_{cat}$ ratio (G768R/WT) | $K_m$ (μM) | $k_{cat}$ ($s^{-1}$) | $k_{cat}$ ratio (G768R/WT) | $K_m$ (μM) | $k_{cat}$ ($s^{-1}$) | $k_{cat}$ ratio (G768R/WT) | $K_m$ (μM) |
| WT (1) | 5.5 | 0.73 | 36.28 | 4.9 | 0.68 | 16.73 | 2.35 | 1.2 | 15.31 |
| G768R (1) | 4.0 | | 23.35 | 3.3 | | 8.09 | 2.8 | | 13.01 |
| WT (2) | 5.3 | 0.69 | 43.57 | 6.3 | 0.72 | 15.91 | 3.3 | 1.15 | 18.64 |
| G768R (2) | 3.67 | | 27.7 | 4.6 | | 9.4 | 3.79 | | 15.89 |
| WT (3) | 5.33 | 0.76 | 36 | 5.8 | 0.58 | 17.01 | 3.24 | 1.2 | 25.83 |
| G768R (3) | 4.1 | | 24.78 | 3.4 | | 8.31 | 3.89 | | 24.33 |
| WT | 5.38 ± 0.06 | | 38.6 ± 2.5 | 5.67 ± 0.41 | | 16.6 ± 0.3 | 2.96 ± 0.31 | | 19.9 ± 3.1 |
| G768R | 3.92 ± 0.13 | 0.73 | 25.3 ± 1.3 | 3.77 ± 0.42 | 0.66 | 8.6 ± 0.4 | 3.49 ± 0.35 | 1.18 | 17.7 ± 3.4 |
| p val | 0.0065 | | 0.010 | 0.017 | | 0.0081 | 0.013 | | 0.027 |

**Table S1.** Fitted $k_{cat}$ and $K_m$ of actin-activated ATPase activity for three different protein preparations (each with 3 technical replicates) of 2-hep, 25-hep and sS1. The respective $k_{cat}$ ratios of G768R/WT for each of the different constructs are shown. The numbers (1), (2) and (3) represent the three different biological replicates. Mean and SEM across the three biological replicates of each protein is shown at bottom rows. p values comparing G768R to WT using the paired t-test are shown in the bottom row. These individual and averaged $k_{cat}$ and $K_m$ values are also plotted in Figure S1.

### Supplementary figures

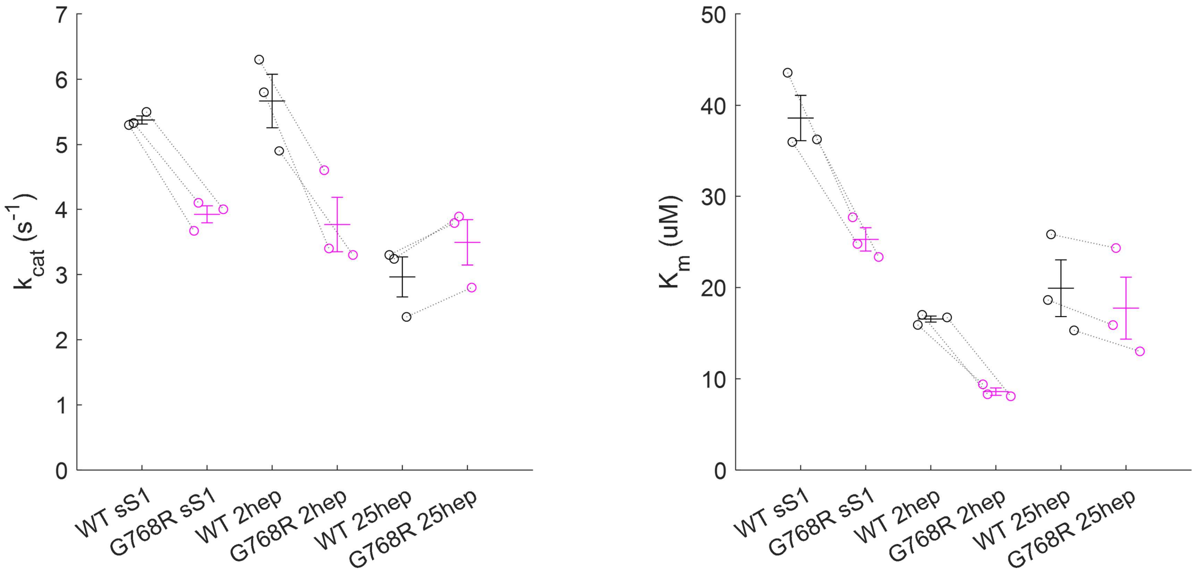


**Figure S1. Actin-activated ATPase of WT and G768R human β-cardiac myosin sS1, 2hep-, and 25hep- HMM.** Data points are fitted $k_{cat}$ and $K_m$ to biological replicates (3 independent paired WT-mutant protein preps for each length construct), with the dotted lines connecting the WT-mutant pairs. Horizontal lines and errors bars represent mean and SEM. All comparisons between G768R and WT are statistically significant with $p<0.05$ using the paired (one-sample) t-test. See Table S1 for numerical values.

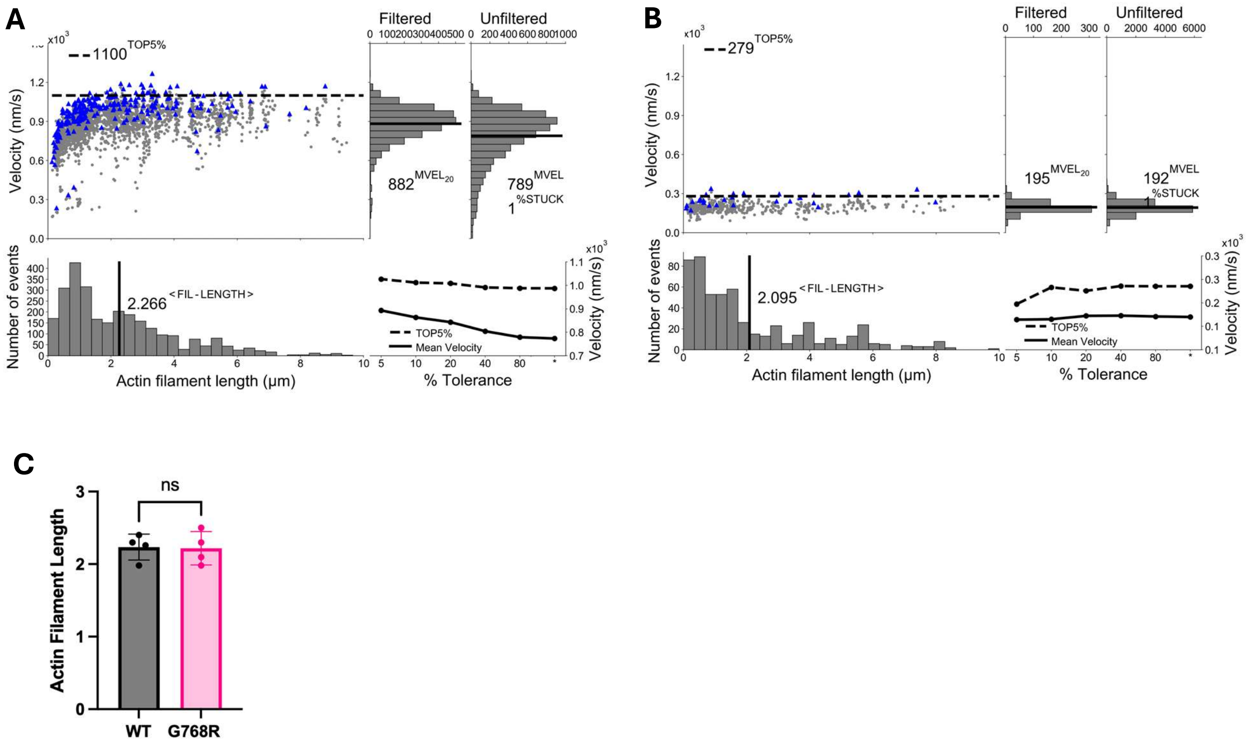


**Figure S2. Representative Fast Automated Spud Trekker (FAST) analysis of in-vitro actin motility data.** The figure shows FAST analysis of actin filament motility data comparing wild-type (WT) **(A)** and G768R mutant **(B)** sS1 proteins. In the top-left scatter plots, each grey point represents the n-frame averaged velocity of a single filament plotted against its length, with multiple points per filament corresponding to different observation frames. Blue points show filtered velocities after excluding filaments with high velocity fluctuations (standard deviation ≥20% of mean velocity when pt=20). The 'Top 5%' dashed line indicates the mean of the highest 5% of filtered velocities, representing filaments with the smoothest, fastest motion. Accompanying histograms display the distributions of filament lengths, filtered velocities, and unfiltered velocities, with solid black lines marking the mean values. The %STUCK value indicates the percentage of essentially immobile filaments (n-frame averaged velocity set to zero when the path-averaged velocity fell below the minimum threshold, minv), which are excluded from the plots. The bottom-right panel shows how filtered mean velocity (MVELpt) and Top 5% values change across different pt (percent tolerance threshold) values, demonstrating the robustness of the analysis parameters. No changes in actin filament length were observed during actin motility assay as shown in **(C)** where each data point represents a biological replicate.

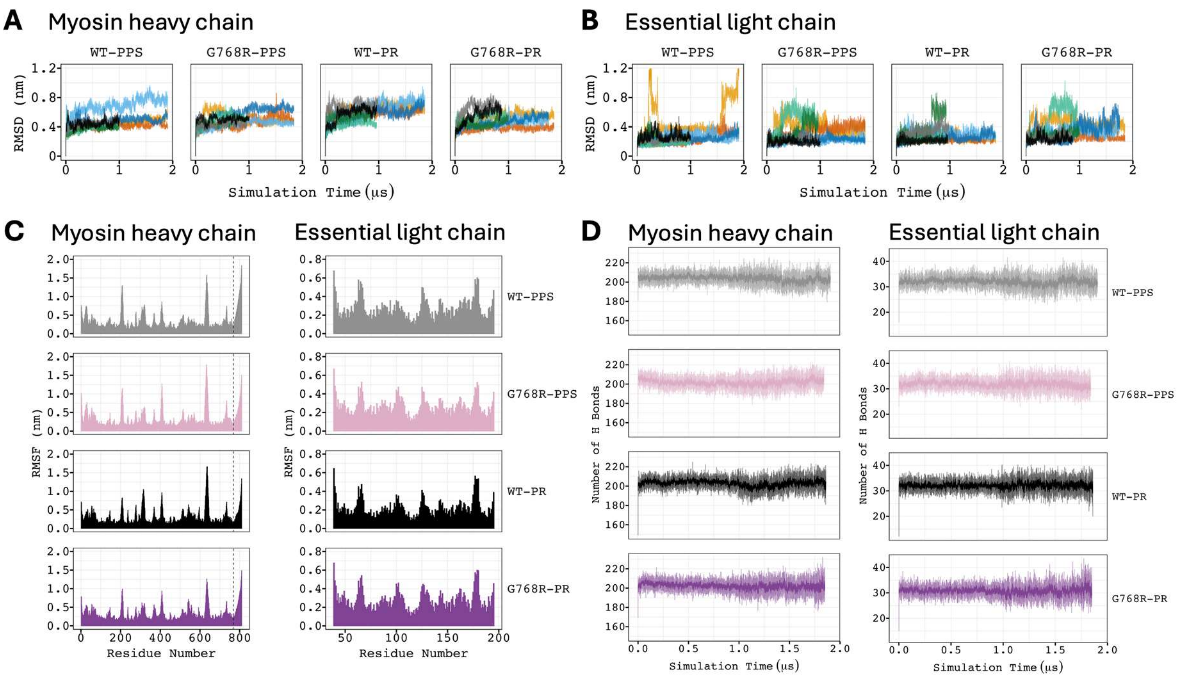


**Figure S3. Structural analysis of WT and G768R myosin models.** RMSD was calculated throughout the separate simulation trajectories of **(A)** all myosin constructs **(B)** their essential light chains. All alpha carbon atoms were used both for the alignment with respect to the initial structural model and for the RMSD calculation. For each construct, the RMSD analysis from 8 independent simulations are colored black, gray, dark blue, light blue, dark green, light green, orange and yellow. **(C)** The root mean square fluctuations per residue was calculated for each construct for myosin (left) and light chain (right) separately using all non-hydrogen atoms of each residue and averaged over all independent simulations. The vertical dashed line shows the residue 768 which is mutated to arginine in the G768R PPS and PR constructs. **(D)** The total number of unique hydrogen bonds was calculated for myosin (left) and light chain (right), separately, as an average over the course of all independent simulations. The solid lines show the rolling average over 100 ps blocks of the trajectories whereas the semi-transparent lines show the raw average. For all subsequent analyses, the first 120-250 ns of each simulation was discarded for further equilibration of the simulation systems.

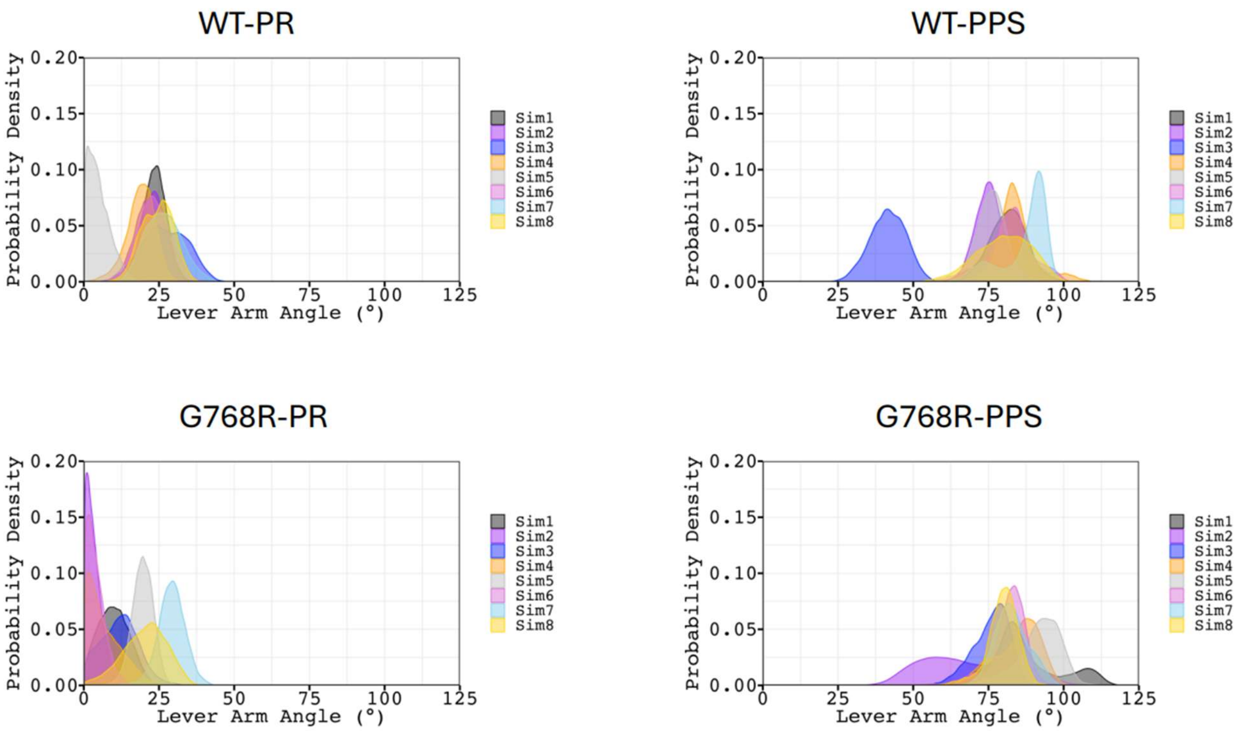


**Figure S4. Distribution of lever arm angle for all individual simulations.** Eight independent simulations of 1-2 μs each were performed for each of the four starting models. Lever arm angle was calculated as the angle of the lever arm in a simulation frame relative to the lever arm of the post-rigor crystal structure (see detailed methods).

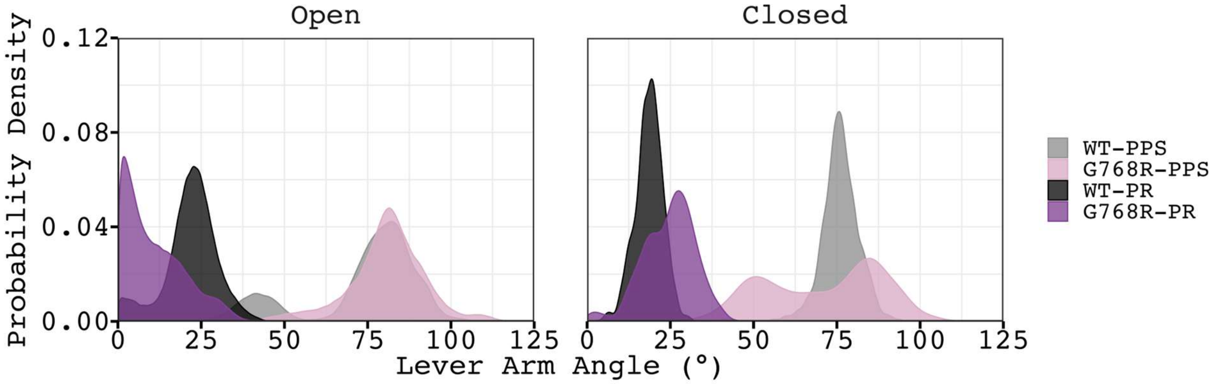


**Figure S5. Distribution of lever arm angle when the inner cleft is open or closed.** The lever arm angle was calculated when the inner cleft distance was open (right) and closed (left). The inner cleft distance was computed as the distance between the alpha carbon atoms of L277 and S472. WT-PPS, G768R-PPS, WT-PR and G768R-PR data are colored gray, pink, black and purple, respectively.

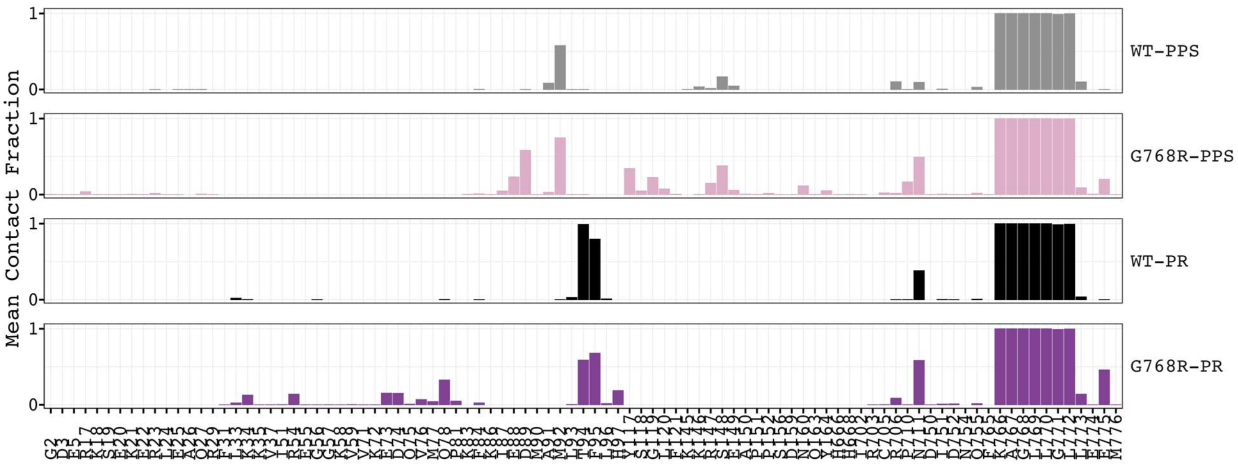


**Figure S6. Myosin contacts with residue 768.** The mean contact fraction for each residue pair was calculated as a time average over 8 independent MD simulations. Residue 768 was considered in contact with a residue if the distance between any non-hydrogen atom pairs between them is less than 0.5 nm.

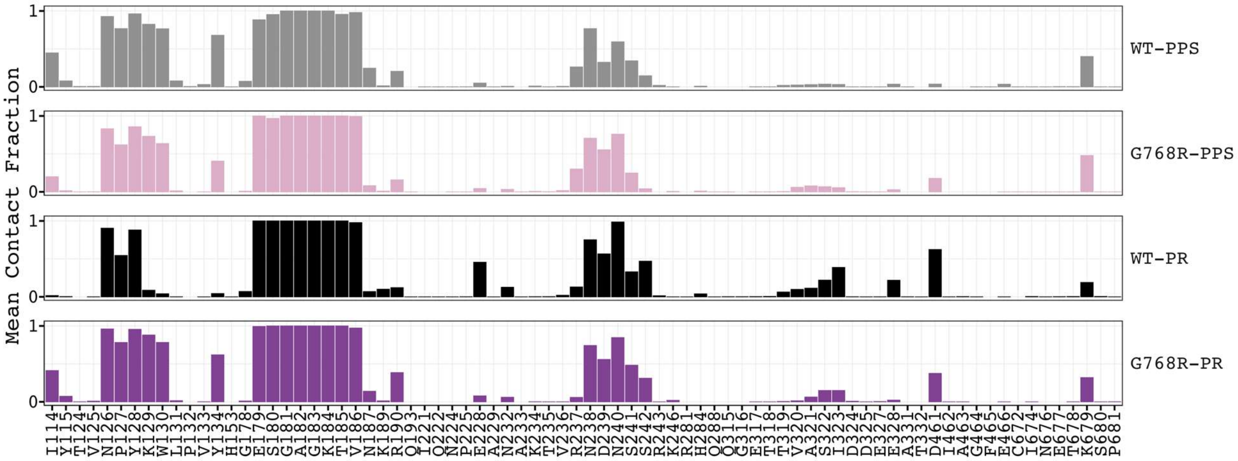


**Figure S7. Myosin-ADP interactions.** The mean contact fraction for each residue-ADP pair was calculated as a time average over 8 independent simulations. ADP was considered in contact with a residue if the distance between any non-hydrogen atom pairs between them is less than 0.5 nm.
